# Evolutionary design of regulatory control. II. Robust error-correcting feedback increases genetic and phenotypic variability

**DOI:** 10.1101/405456

**Authors:** Steven A. Frank

**Author notes:** web: https://stevefrank.org.

## Abstract

As systems become more robust against perturbations, they can compensate for greater sloppiness in the performance of their components. That robust compensation reduces the force of natural selection on the system’s components, leading to component decay. The paradoxical coupling of robustness and decay predicts that robust systems evolve cheaper, lower performing components, which accumulate greater mutational genetic variability and which have greater phenotypic stochasticity in trait expression. Previous work noted the paradox of robustness. However, no general theory for the evolutionary dynamics of system robustness and component decay has been developed. This article takes a first step by linking engineering control theory with the genetic theory of evolutionary dynamics. Control theory emphasizes error-correcting feedback as the single greatest principle in robust system design. Linking control theory to evolution leads to a theory for the evolutionary dynamics of error-correcting feedback, a unifying approach for the evolutionary analysis of robust systems. In this article, I study how, in theory, increasingly robust systems accumulate more genetic variability and greater stochasticity of expression in their components. The theory predicts different levels of variability between different regulatory control architectures and different levels of variability between different components within a particular regulatory control system. Those predictions provide a way to understand the accumulating data on genetic variability and single-cell stochasticity of gene expression. I also show that increasing robustness reduces the frequency of system failures associated with disease and, simultaneously, causes a strong increase in the heritability of disease. Thus, robust error correction in biological regulatory control may partly explain the puzzlingly high heritability of disease and, more generally, the surprisingly high heritability of fitness.

## Introduction

As a system’s robustness increases, it becomes less sensitive to perturbations. A robust system also becomes less sensitive to the effects of deleterious mutation, a particular kind of perturbation. Less sensitivity to mutation means that the force of natural selection against mutation becomes weaker. That weakened selection leads to the accumulation of more mutations and increased genetic variability, a consequence of *mutational robustness* (Rutherford & Lindquist, 1998; de Visser et al., 2003; Wagner, 2013). Previously, I suggested a more general principle that influences the design dynamics of all systems, *the paradox of robustness.* The better a system becomes at compensating for perturbations and errors, the more the system’s components will tend to decay in performance (Frank, 2004, 2007b, 2013). Better error correction begets more errors.

Component decay may take the form of increased variability or sloppiness in function, of which mutational variance is one special case. Alternatively, when robust systems can compensate for poor components, the system’s components may decay to less costly and lower performing types. In that case, the economics of efficiency favors robust systems to use cheaper components.

Such economic arguments of efficiency seemingly must apply to the design dynamics that shape all systems. However, no broad theory has developed general aspects of robust system design dynamics in relation to the decay of system components. How does the ongoing duality of improved system robustness and component decay shape the path by which complex systems are created? What are the consequences for system characteristics?

This article takes an initial step toward a broad theory. I focus on error-correcting feedback for the initial analysis of robustness, because error correction is the single greatest principle of robust system design (Åström & Murray, 2008; Ogata, 2009; Dorf & Bishop, 2016; Frank, 2018a).

In an error-correcting feedback system, the error measures the difference between a system’s actual output and its target. By feeding back the error as an input, the system can move in the direction that reduces the error. Error correction compensates robustly for misinformation about system dynamics and for perturbations to system components. Excellent performance often follows in spite of limited information, sloppy components, and noisy signals.

Engineering control theory provides a rich, highly developed theory of error-correcting feedback. I connect the insights of control theory with the well developed theory of evolutionary dynamics for genetic systems. This link between control theory and genetics provides a first step toward a theory for the evolutionary dynamics of feedback control.

This article focuses on the consequences of robust feedback control for the patterns of genetic and phenotypic variability that arise by evolutionary dynamics. The final *Conclusions* section summarizes key results and promising directions for future study.

## Background

The first article in this series developed the methods to analyze performance, dynamics, and feedback loops (Frank, 2018b), based on the general principles of control theory (Åström & Murray, 2008; Ogata, 2009; Dorf & Bishop, 2016; Frank, 2018a). This section briefly reviews the methods of that first article. I will then use those methods to develop new analyses of genetic and phenotypic variability in relation to control architecture.

### Performance and fitness

I focus on two components of performance, homeostasis and tracking. Those two components typically trade off. A good homeostatic system tends to respond weakly to external disturbance inputs but, in consequence, tends to be a poor tracking system that only slowly follows the long-term changes in environmental inputs.

For homeostasis, I analyze the system’s response to a single intense perturbation of short duration. Performance measures the deviation of the system from its homeostatic setpoint

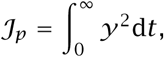

in which *y(t)* is the system output at time *t*, and *y =* 0 is the target setpoint for the system. Thus, *𝒥*_*p*_ measures the total squared deviation of the system in response to a perturbation. I use the classic Dirac delta impulse perturbation applied at time zero, which imposes an input of infinite intensity and infinitesimal duration at time zero.

For tracking, I analyze the error deviation, *e = y - u*, in which each term depends on time, *t*, with *u* as the environmental input that sets the system target state, yielding the measure of tracking performance

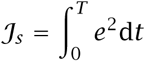

from an initial time *t =* 0 until some final time, *t = T.* I will typically use *T =* 20, which is sufficient to describe how well a system adjusts to an environmental change. The error deviations are measured in response to a step-change in the environmental input, *u(t) =* 0 for *t <* 0 and *u(t) =* 1 for *t ≥* 0. This tracking component of performance, *𝒥*_*s*_, captures the system’s step response.

Total performance is a combination of step and perturbation response

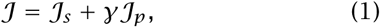

in which *γ* weights the importance of the homeostatic perturbation response relative to the tracking step response.

Lower values of *𝒥* correspond to smaller squared deviations from target phenotypes. Thus, minimal values of *𝒥* correspond to maximal performance.

For evolutionary analysis, we can transform performance measures into fitness as

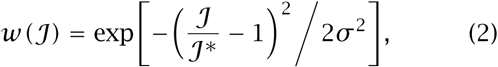

in which *𝒥\** is the optimum (minimum) value of the performance measure, thus optimum (maximum) fitness is one, and minimum fitness is zero. Smaller values of *σ* ^2^ correspond to a more rapid decline in fitness with a change in performance. I typically use *σ*^2^ *=* 0.01, which corresponds to *σ =* 0.1, and approximately a 40% decline in fitness for a 10% reduction in performance.

### Dynamics and transfer functions

We can transform the temporal dynamics in the time variable *t* for a differential equation such as

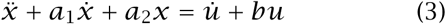

into an expression in the complex Laplace variable *s* as

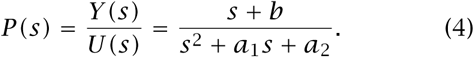

The numerator expresses a polynomial in *s* derived from the coefficients of *u* from the right side of eqn 3. Similarly, the denominator expresses a polynomial in *s* derived from the coefficients in *x* from the left side of eqn 3. The eigenvalues for the process, *P*, are the roots of *s* for the polynomial in the denominator.

From eqn 4, we may write *Y* (*s*) = *U*(*s*)*P*(*s*). In words, the output signal, *Y* (*s*), is the input signal, *U*(*s*), multiplied by the transformation of the input signal by the process, *P* (*s*). Because *P* (*s*) multiplies the signal, we may think of *P* (*s*) as the signal gain or amplification, which is the ratio of output to input, *Y* /*U*.

The simple multiplication of the signal by a process means that we can easily cascade multiple input-output processes. In Fig. 1a, a preprocessing controller of the form *C*(*s*) = *U*(*s*)/*R*(*s*) takes an external environmental input signal, *R*, and outputs a control signal, *U*. Thus, we can write a cascade, *R* → *C* → *P* → *Y*, that takes input, *R*, and transforms that input via a preprocessing controller, *C*, and a process, *P*, to yield an output, *Y*, as

**Figure 1:**
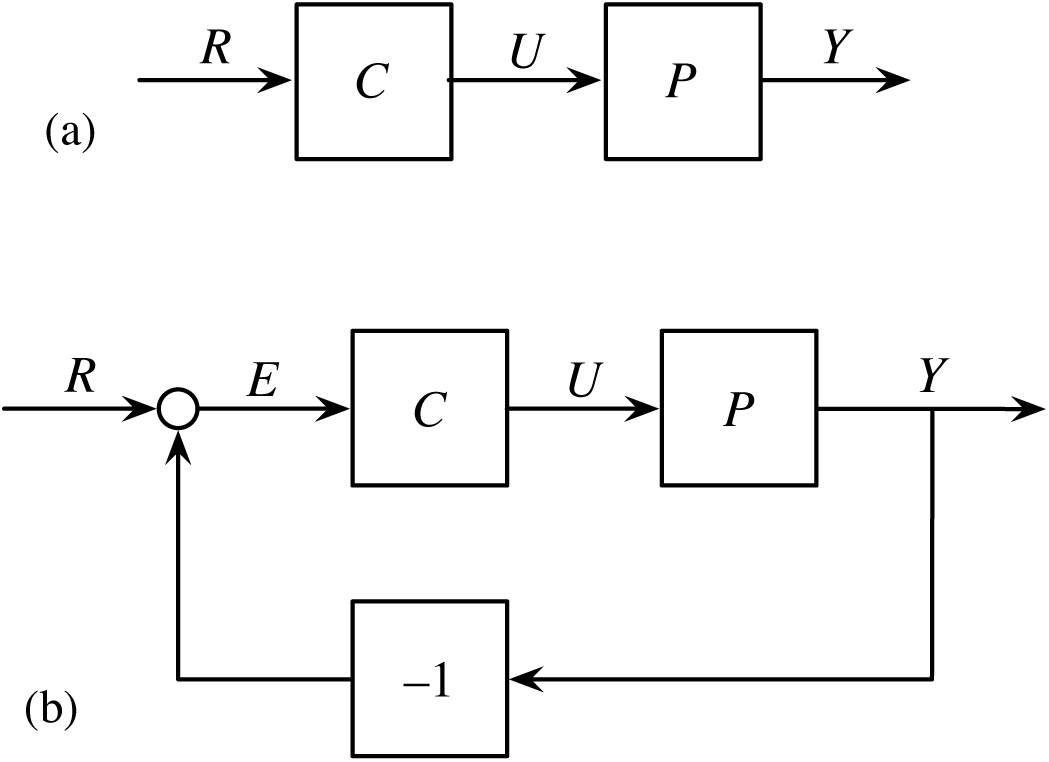
Control systems in which a modifiable controller, *C*, alters the dynamics of an intrinsic, unmodifiable plant process, *P.* (a) Open loop system, for which the input signal, *R*, leads to the output, *Y*, through *R* → *G* → *Y* for the internal open loop processing system, *G* = *CP*. (b) Closed feedback loop, for which the input signal, *R*, leads to the output, *Y*, through *R* → *G* → *Y* for the internal closed loop processing system, *G* = *CP*/(1 + *CP*), as described in eqn 5.

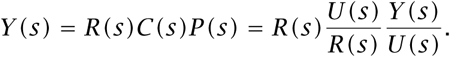

These functions of *s* are called *transfer functions*.

### Feedback loops

With transfer functions, we can easily calculate the total system response of a feedback loop. Consider the steps by which we can analyze the dynamics of the feedback loop in Fig. 1b.

If we write all signals and internal processes as transfer functions, with *Y* as the transfer function for the output signal, *R* as the transfer function for a system setpoint given as the system input, and *E* = *R* − *Y* as the transfer function for the error between the external setpoint and the actual output, then the direct line of signal processing between the input and the output without feedback yields an output *Y = CPE*, because transfer functions multiply along a signal line. Substituting *E = R - Y* into the previous input-output expression, we obtain

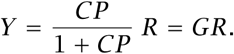

The complete feedback loop system, *G*, that takes input *R* and yields output *Y* is

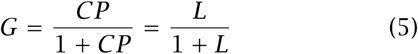

in which *L = CP* is often called the open loop component of the system—the open part of the system without the feedback that closes the loop.

### Optimized loop example

The analyses in this article begin with simple control loop examples, with structures such as those in Fig. 1. I use the intrinsic, unmodifiable “plant” process

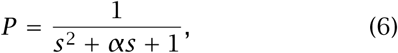

which corresponds to the dynamics of the second-order differential equation

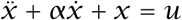

for system state *𝓍 (t)* and input *u(t)*, with the single parameter *α*, and with system output equivalent to system state, *y* ≡ *𝓍*. Optimizing the performance measure in eqn 1 yields 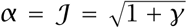. I use this optimal value for *α* in all analyses. Thus, we can study how modifiable processes can b e added t o a system to achieve improved performance.

For the open loop in Fig. 1a, the controller, *C*, alters the input signal that is passed into the intrinsic process, *P.* For the controller, I use

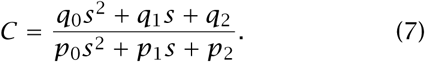

Optimization methods find the best values of the *q*_*i*_ and *p*_*i*_ parameters. With optimized parameters, we can then study the sensitivity of the system to genetic or phenotypic variability in those parameters. Numerical optimization (Frank, 2018b) based on the performance measure in eqn 1 suggests that the optimal controller typically transforms the uncontrolled second order plant system, *P*, into the first order open loop controlled system *G = CP* in Fig. 1a, in which

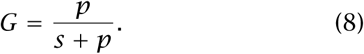

The optimized controller is

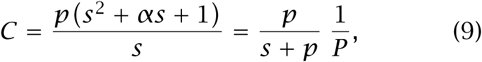

with 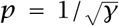. The optimized open loop has performance 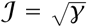, which improves on the optimized unmodified plant, *P*, with performance 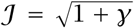. The optimized parameters of *C* with respect to the general form in eqn 7 are

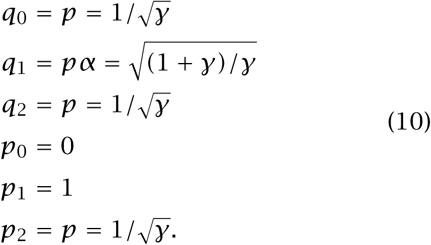

Figure 1 shows that both open and closed loops process the input signal, *R*, into the output signal, *Y*, through some internal signal processing, *G*, such that the systems can be written as *R* → *G* → *Y.* Thus, for optimized open and closed loops, *G* typically takes the same form, even though the two control loop architectures arrive at that final overall processing by different controllers.

For the open loop, *G = CP*, and for the closed loop, *G* is given in eqn 5. For the closed loop to produce the same optimal *G* as the open loop, given in eqn 8, the optimal closed loop controller is

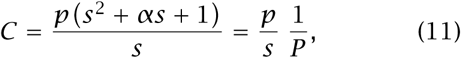

with parameters equivalent to those in eqn 10, except that *p*_2_ *=* 0.

## Sensitivity of control loops

To study the sensitivity of systems to genetic and phenotypic variation, we can analyze changes in performance with changes in the controller parameters given in the previous section. For the examples in this section, I use *γ =* 2.

### Open vs closed loop sensitivity

Figure 2a compares the sensitivity of the open versus closed loops in Fig. 1 for changes in the controller parameters. For the open loop (blue curves), I initially set all parameters to the optimum values in eqn 10. Then, for each parameter one at a time, I varied the value over *θ*2^*x*^ for the optimal parameter value, *θ*, and the range of *x* over ±2.

**Figure 2:**
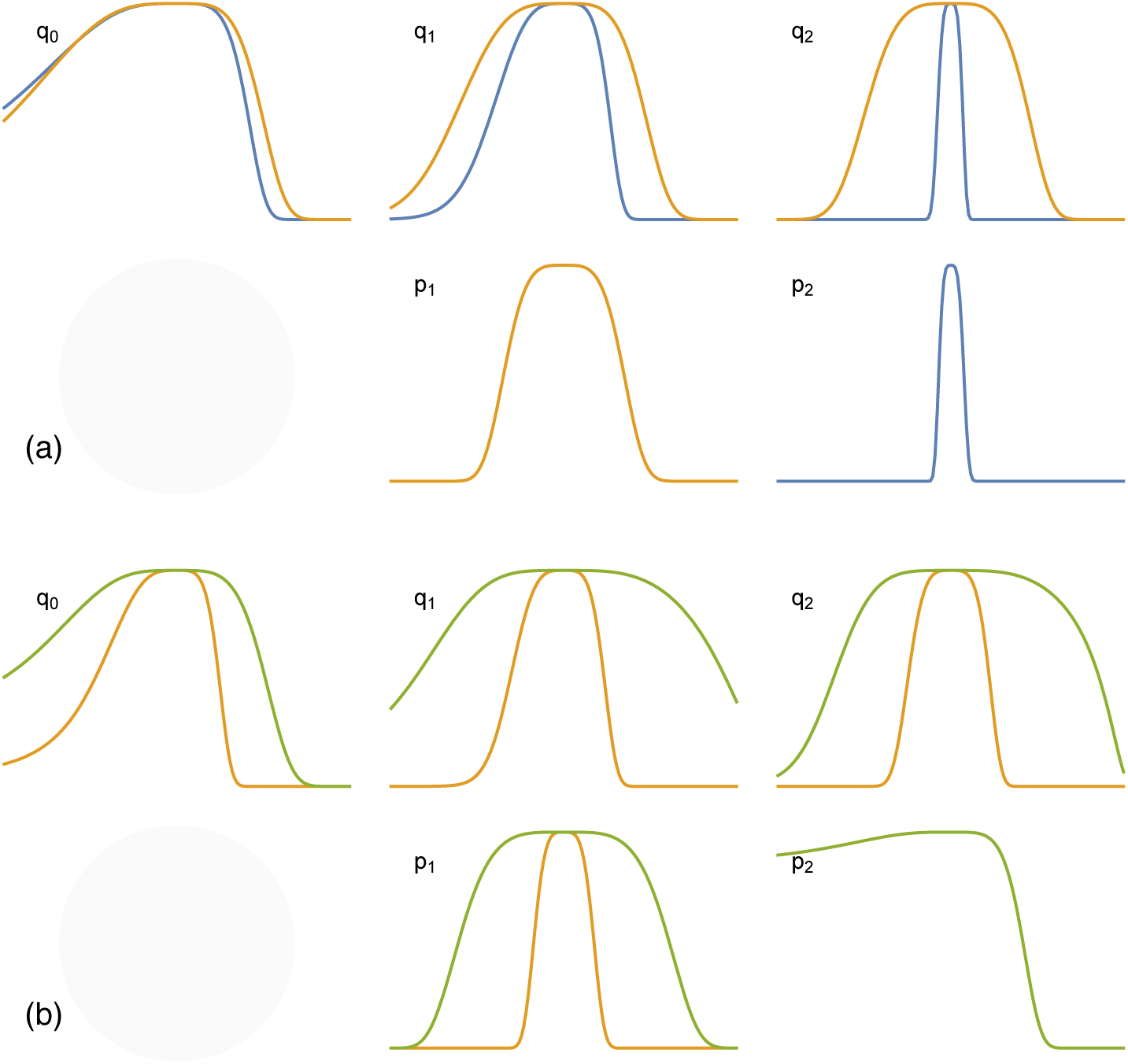
Sensitivity comparison between systems. (a) Open loops (blue curves) versus closed loops (gold curves) with respect to variations in the controller parameters. The *x*-axis covers *θ*2^*x*^, with *θ* as the optimal value and *x* over the range ±2. The height of each curve is the fitness relative to a maximum value at the optimal parameter value in the center of each plot at *x =* 0. The optimal parameter for *p*_0_ is zero, thus that parameter is not shown. The optimal closed loop parameter for *p*_2_ is zero, thus there is no gold curve for the closed loop for that parameter. For *p*_1_, the sensitivity curves overlap, thus only the gold curve appears. (b) Single closed loop (gold curves) versus double closed loop (green curves). The *x*-axis range is ±4. The single loop gold curves match the gold curves from the upper panels, adjusted for the different *x*-axis scaling.

For each parametric variation, I calculated the performance measure in eqn 1, and then transformed performance into fitness in eqn 2. Thus, each curve shows the variation in fitness as a function of variation in a single parameter, holding the other parameters constant at their optimal values.

For the closed loop (gold curves), I followed the same procedure. The only difference is that the optimal closed loop value for the parameter *p*_2_ is zero, thus there is no closed loop gold curve for that parameter.

The closed loop is less sensitive to variations in the *q* parameters. The intrinsic error correction of feedback compensates for parametric variations when compared to an open loop, which lacks any intrinsic correction for errors. The closed loop *q* parameters would be expected to maintain significantly greater genetic variability.

For a given genotype, phenotypic fluctuations in the less sensitive closed loop parameters would have less fitness consequence than similar fluctuations in open loop parameters. Thus, natural selection for regulation of expression would be weaker, and greater observed phenotypic stochasticity in those parameters would be expected.

The relative relaxation of sensitivity for the various *q* parameters differs, leading to predicted differences in genetic and phenotypic variability. This example suggests that one may construct a fundamental theory for genetic and phenotypic variability in relation to the architecture of control and the associated sensitivities of particular system components.

The error correction of feedback loops may create particularly strong robustness and insensitivity to parametric fluctuations. Thus, a natural association arises between enhanced system robustness and associated genetic and phenotypic decay in the performance of certain system components (Frank, 2004, 2007b, 2013).

### Double closed loop sensitivity

Robustness mechanisms in the form of feedback loops may be layered on top of each other in control architectures. For example, Fig. 3 shows a double feedback loop. The inner loop matches the basic feedback architecture of Fig. 1b. The outer loop adds another error correcting layer of control. How much does the extra robustness of the outer loop reduce the sensitivity of the controller parameters of the inner loop?

**Figure 3:**
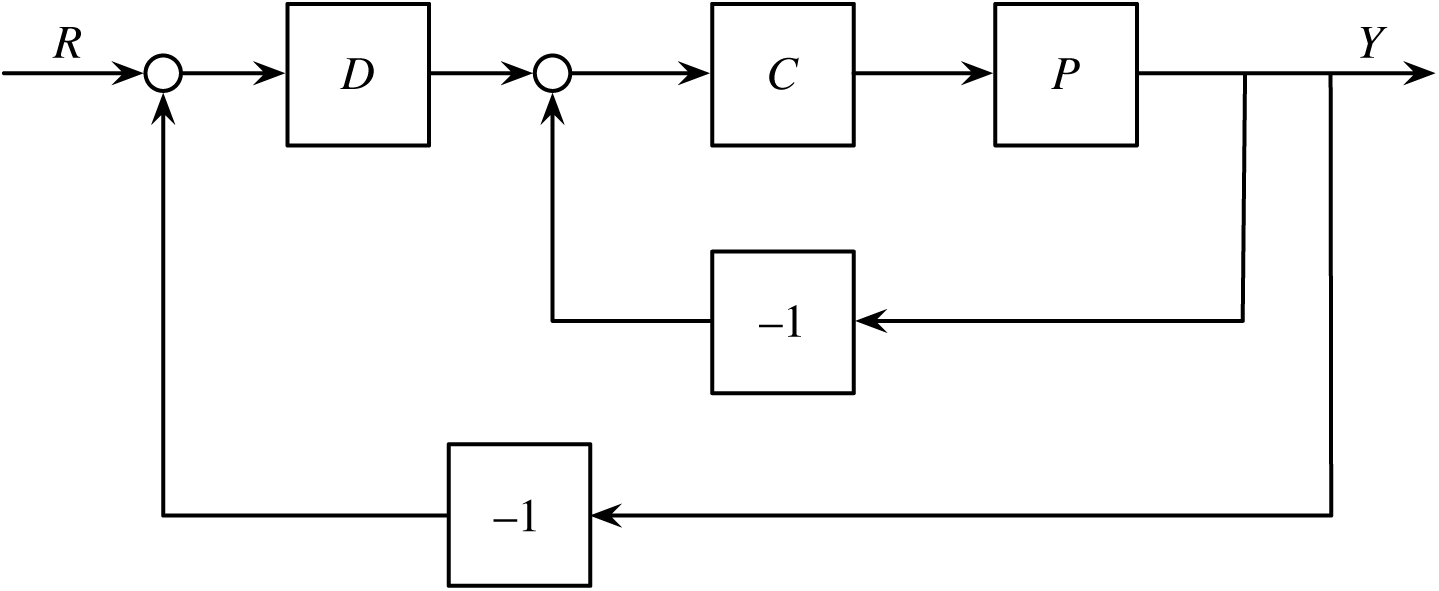
Double closed loop feedback. The inner loop is identical to the single closed loop in Fig. 1b. The outer loop provides a second error-correcting mechanism that potentially reduces the system’s sensitivity to parameter variations in the inner loop.

Figure 2b compares the sensitivities of a single closed loop (Fig. 1b) in the gold curves with the sensitivities of the double closed loop (Fig. 3) in the green curves. The double loop greatly reduces the sensitivities of the controller parameters. That enhanced robustness and reduced sensitivity of the double loop will lead to the accumulation of greater genetic variability and the expression of greater phenotypic stochasticity.

To calculate the double loop sensitivity, one has to start with the altered optimal parameters for the controller, *C*, and the optimal parameters for the additional system component, *D*. We begin with the inner loop given earlier: *L = CP* and *G = L/(*1 *+ L)*. Then, applying the same logic to the outer loop, we can write the complete system as *H* = *DG*/(1 *+ DG*). The optimized full system, *H*, has the same dynamics as the previous optimized systems, *H* = *p*/(*s* + *p*) with 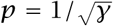.

Let the parametric form of the outer controller be *D = k + kr/s*, a standard proportional-integral controller from basic control theory (Åström & Murray, 2008; Ogata, 2009; Dorf & Bishop, 2016; Frank, 2018a). Then the optimal controller is

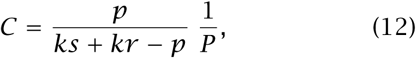

which yields the controller parameters as in eqn 10 with *p*_1_ *= k* and *p*_2_ *= kr - p*. In the numerical example shown in Fig. 2b, I used *r =* 10 and *k =* 1*/r.*

Figure 4 shows the effects of varying *r* or *k* on aspects of system sensitivity. In Fig. 4a, I varied *r* or *k* within the outer loop controller, *D*, but kept the default values of *r =* 10 and *k =* 1*/r* for the inner controller, *C*. The plot shows that system performance does not change significantly when *r* or *k* is multiplied by a factor of 2^*x*^ for *x* ≈ 1/3, which means altering the parameter by approximately ±26%.

**Figure 4:**
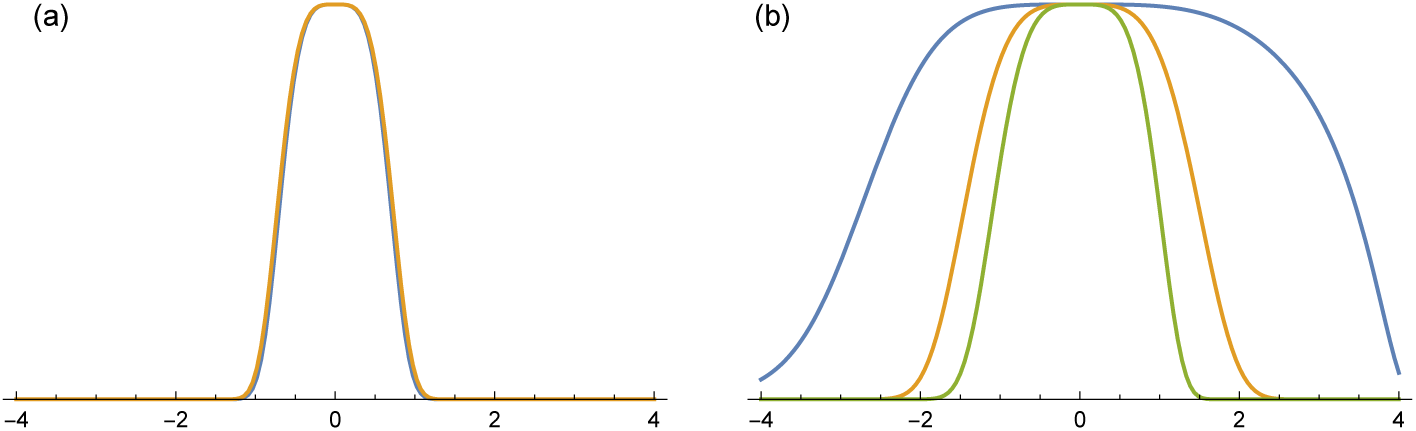
Sensitivity of the double feedback loop to variations in the additional *r* and *k* parameters introduced with the double loop architecture. The *x*-axis is the value of the parameter with the optimum at zero and varying over the range 2^*x*^ for *x* ± 4. (a) Sensitivity of fitness to changes in *k* (blue) and *r* (gold) of the outer controller, *D*, holding the inner loop values of those parameters in the controller *C* at their optimum. The overlapping curves are approximately the same. (b) Sensitivity of the inner loop controller parameter, *q*_2_, for different values of *k* in both the inner and outer loop controllers, with *k =* 1*/r* (blue), *k =* 2*/r* (gold), and *k =* 8*/r* (green).

Figure 4b shows the effect of varying *k* in both the inner and outer loops on the sensitivity of the parameter *q*_2_ of the controller, *C*. The outer blue curve of Fig. 4b corresponds to *k =* 1*/r*, which matches the outer green curve of Fig. 2b for *q*_2_. The middle gold curve of Fig. 4b corresponds to *k =* 2*/r.* The inner green curve of Fig. 2b corresponds to *k =* 8*/r*, which approximately matches the inner gold curve of Fig. 2b. Thus, the reduced sensitivity of the inner loop parameters induced by the double feedback loop architecture is itself sensitive to variation in the matching *k* parameter of the inner and outer loops.

## Computer simulations

The previous sections suggested that additional error-correcting feedback enhances robustness and reduces sensitivity. To evaluate those predictions, I studied an evolutionary model of the various control loop designs with respect to fitness and aspects of genetic and phenotypic variability.

I used computer simulations to handle the full complexity of the genetics, expression of traits, and calculation of fitness. The Supplementary Information (SI) describes the details of the simulations and the parameters used in various simulation experiments. The individual simulation experiments vary subsets of parameters factorially, as described in the SI. The SI also provides additional analyses of the simulation output. The following sections highlight the key results.

## Robustness and fitness

How does the robustness of error-correcting feedback influence fitness? To study that question, we can compare fitness for the three control loop designs: open loop, single error-correcting feedback loop, and double error-correcting feedback loop. In theory, greater error correction should enhance robustness. We can analyze robustness by studying the mean and standard deviation in fitness for evolving populations with respect to varying levels of perturbation to the intrinsic plant process, *P.*

Figure 5 shows that increased error correction robustly protects against perturbations. For example, the green line in Fig. 5a shows the mean fitness response of the three control designs when the plant parameter, *α*, in eqn 6 is strongly perturbed. The error correction of the single (S) feedback loop significantly increases fitness compared to the open (O) loop that cannot correct errors. The double (D) feedback loop can restore the same high fitness level as in the case for which the plant parameter is not perturbed (blue line).

**Figure 5:**
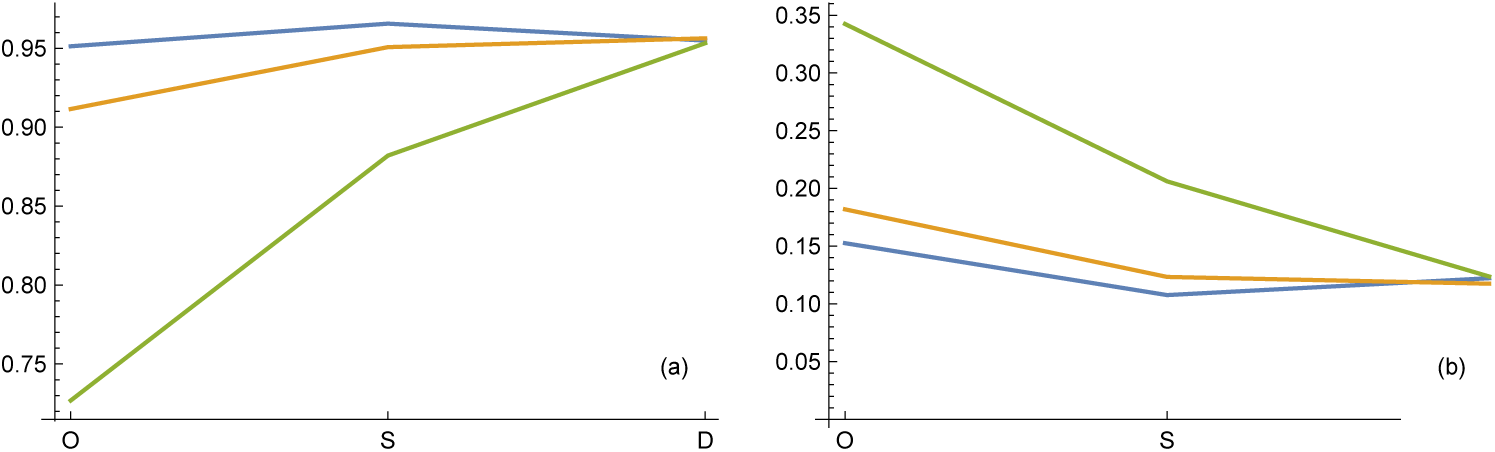
Plots of (a) mean fitness and (b) standard deviation in fitness in relation to open (O), single (S), and double (D) control loops. The lines show different levels of perturbation to the intrinsic plant parameter in eqn 6, 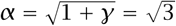, for *γ =* 2. The value of *α* in an individual is 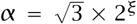, in which ξ is a random number drawn from a Gaussian distribution with mean of zero and standard deviation of *aSD*. The values of *aSD* are 0 (blue lines), 0.25 (gold lines), and 0.5 (green lines). Results from Experiment A as described in the Supplementary Information.

Similarly, in Fig. 5b, the variation in fitness in the population for the double (D) loop remains no greater under strong perturbation to the plant (green line) than for the unperturbed case (blue line). In summary, single error correction provides some protection against perturbation and lessens the effects of perturbation, whereas double error correction provides essentially perfect protection against perturbation.

The simulations summarized in Fig. 5 also have mutation, which causes variation in the genetically determined component of the controller parameters, and nongenetic stochastic fluctuations in the controller parameters (see SI). Those fluctuations in controller parameters prevent the population from reaching an optimal mean fitness of one. However, those controller fluctuations do not cause significant differences in the success of the three control loop designs, as can be seen in the relative flatness of the blue lines.

Figure 6 shows the relative robustness of the three control loop designs with respect to two other parameters. In the top panels (a,b), the lines from green to blue show an increasing rate of change in fitness with changes in performance, corresponding to a decline in *σ* ^2^ in eqn 2. In other words, the blue lines show high sensitivity of fitness to small changes in performance. High sensitivity would occur if the performance characteristics had a strong influence on the organism’s success. By contrast, the weaker sensitivity of fitness to performance in the green lines would occur if performance for the particular trait played a smaller overall role in the lifetime success of an organism.

**Figure 6:**
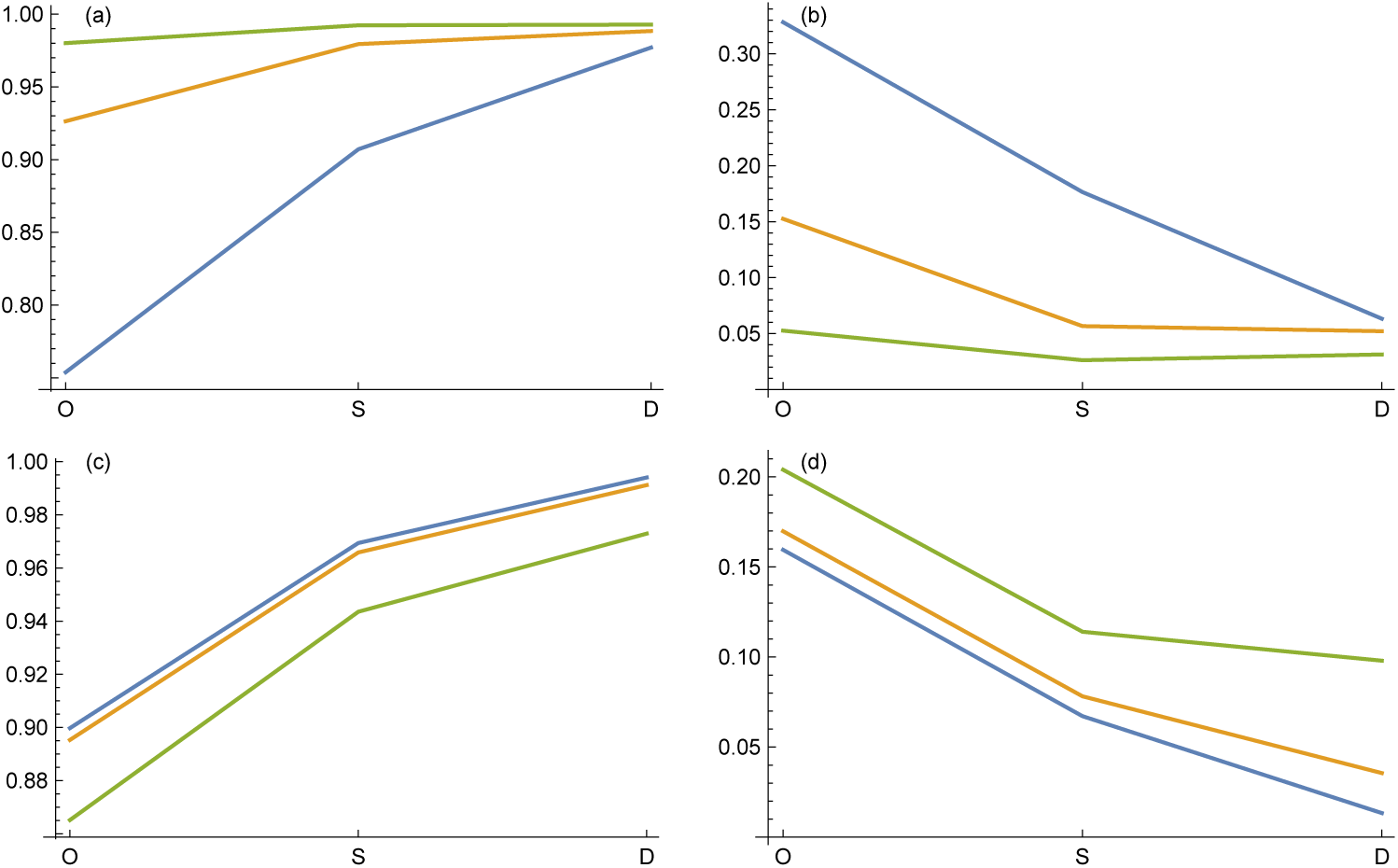
Plots of (a,c) mean fitness and (b,d) standard deviation in fitness, as in Fig. 5. (a,b) The lines show different levels of *σ* ^2^, the parameter in eqn 2 that determines the rate of change in fitness, *𝒲*(*𝒥*), with change in performance, *𝒥*. The values of *σ* ^2^ are 0.01 for relatively rapid change in fitness with change in performance (blue lines), 0.1 for relatively intermediate change in fitness with change in performance (gold lines), and 1 for relatively slow change in fitness with change in performance (green lines). (c,d) The lines show different levels of the mutation rate, with 10^*-*4^ (blue lines), 10^*-*3^ (gold lines), and 10^*-*2^ (green lines). Results from Experiment B as described in the Supplementary Information.

The results in Fig. 6a,b show, once again, that a single error-correcting loop provides significantly improved robustness relative to an open loop. The double loop achieves almost complete robust protection, even when fitness becomes very sensitive to small changes in performance (blue lines).

In the bottom panels, Fig. 6c,d, the mutation rate increases from the blue to the green lines. The very high mutation rate in the green lines degrades fitness and increase variability, whereas the lower, more realistic rates in the blue and gold lines show that as mutation rate declines, fitness mean and standard deviation change relatively little.

## Genetic and phenotypic variability

As the robustness of error-correcting feedback increases, we may expect evolutionary dynamics to accumulate more variability in system parameters. Put another way, better error correction at the system level begets more errors at the component level.

Particular components vary in their tendency to accumulate variability. The tendency to vary depends on how much enhanced error correction reduces the sensitivity of fitness to variability in a particular component. Figure 2 shows how each additional error-correcting feedback layer reduces the sensitivity of particular components.

For example, the sensitivity of fitness to variability in the controller parameter *q*_2_ decreases greatly from the least robust open loop (blue) to the single feedback loop (gold) to the double feedback loop (green). By contrast, enhanced robustness changes fitness sensitivity relatively less with respect to variations in *q*_0_.

In Fig. 2, the analysis of variability for each component assumed that all other components were fixed at their optimum value. In an actual population, all of the components vary simultaneously. Additionally, relative sensitivity provides insight into relative variability, but does not by itself determine the actual amount of variability. The computer simulations provide insight into robustness and component variability that arise by evolutionary dynamics. The Supplementary Information describes the simulation methods and data analysis.

In the simulated populations, two inherited genes influence the variability of each controller parameter. The expressed parameter value in an individual is the value encoded by its first gene multiplied by 2^*z*^, in which *z* is a random number drawn from a Gaussian distribution with mean of zero and standard deviation of *stochWt × δ*. The value of *stochWt* is a parameter of the simulated population, and *δ* is the value encoded by the second gene. The two genes may be thought of as encoding the inherited genetic value of a parameter and the inherited tendency for phenotypic variability in expression of that parameter.

Figure 7 summarizes the variability of the controller parameters in a simulated population. The top-left panel shows the variability in the inherited genetic value of the parameter *p*_1_ over all individuals in the population. Each curve is the cumulative distribution function (CDF) for the probability distribution of genetic values.

**Figure 7:**
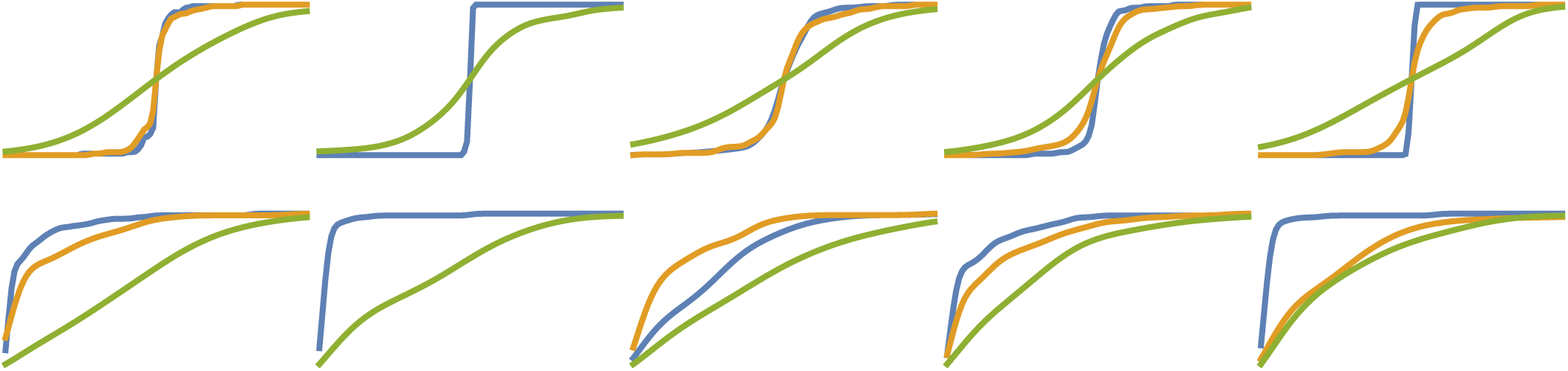
Variability of system components in simulated populations. Each curve is a cumulative distribution function of inherited values. The colors show different control architectures corresponding to different levels of error-correcting feedback and robustness: open loop with no feedback (blue), single error-correcting feedback loop (gold), and double error-correcting feedback loop (green). The columns from left to right show variability for the parameters *p*_1_*, p*_2_*, q*_0_*, q*_1_, and *q*_2_. See text for explanation. In the second column, for the single feedback loop, *p*_2_ has its optimum value at zero, thus the associated gold curve does not appear. The Supplementary Information provides full details of the methods and simulation data.

Along the horizontal axis, the curve is centered at the median genetic value of the distribution, at half the maximum height of the CDF curve, which varies between zero and one. The genetic values along the horizontal axis vary logarithmically from the median multiplied by 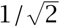 to the median multiplied by 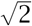. The flatter the CDF curve, the more widely variable the genetic values are in the population. The fre quency of any small interval of genetic values along the horizontal axis is proportional to the slope of the CDF curve.

The three CDF curves in each upper panel show the different control architectures. The blue curve plots an open loop with no error correction, the gold curve plots a single error-correcting feedback loop, and the green curve plots a double error-correcting feedback loop. Robustness increases from the lowest robustness represented by the blue curve to the highest robustness represented by the green curve, with the gold curve in the middle.

The double feedback loop (green), which is the most robust, accumulates the greatest inherited genetic variability in all parameters. The open loop (blue), which is the least robust, accumulates the least amount of variability.

These patterns of variability for the inherited genetic value match the main prediction that increasing system robustness corresponds to greater component variability.

The lower panels show the inherited tendency for stochastic variability in phenotypic expression. The horizontal axis ranges from zero, for no stochasticity, to one, for relatively high stochasticity. CDF curves that are flatter and to the right correspond to greater stochasticity of expression. Again, the tendency is for the most robust double feedback loop to have the greatest variability, which in this case corresponds to higher values of stochasticity.

These results for variability tend to follow the predicted sensitivities in Fig. 2. Thus, we can use fundamental theory to predict how particular components of a regulatory control system accumulate genetic variability and stochastic variability in phenotypic expression. The theory suggests that certain components in the regulatory control system will tend to be highly variable, whereas other components will be less variable.

The Supplementary Information provides details about the simulation methods, the various simulation experiments, the simulation data, and the methods of data reduction.

## Heritability of disease

Many human diseases have a significant heritable genetic component (Frank, 2007a; Manolio et al., 2009; Eichler et al., 2010). Why doesn’t natural selection remove the deleterious genes that cause disease? The paradoxical nature of robustness may partly explain the high heritability of disease (Frank, 2004).

Increased robustness protects against the failures caused by perturbations, reducing disease. Better robustness also allows increased inherited variability in each component of the system (Fig. 7). Thus, when a robust system fails in disease, it may be more likely to do so because an individual has inherited faulty components. In other words, robustness may reduce the frequency of disease and raise the heritability of disease.

Figure 8 shows that robustness does reduce disease frequency and increase the heritability of disease. In that figure, disease is defined as a reduction in fitness of greater than 10% from maximum fitness. The horizontal axes show the percentage of the population that express disease. As robustness increases from open loop (blue) to single feedback loop (gold) to double feedback loop (green), the frequency of disease declines. The symbol *Φ* denotes the frequency of disease, with 100*Φ* as the percentage of disease.

**Figure 8:**
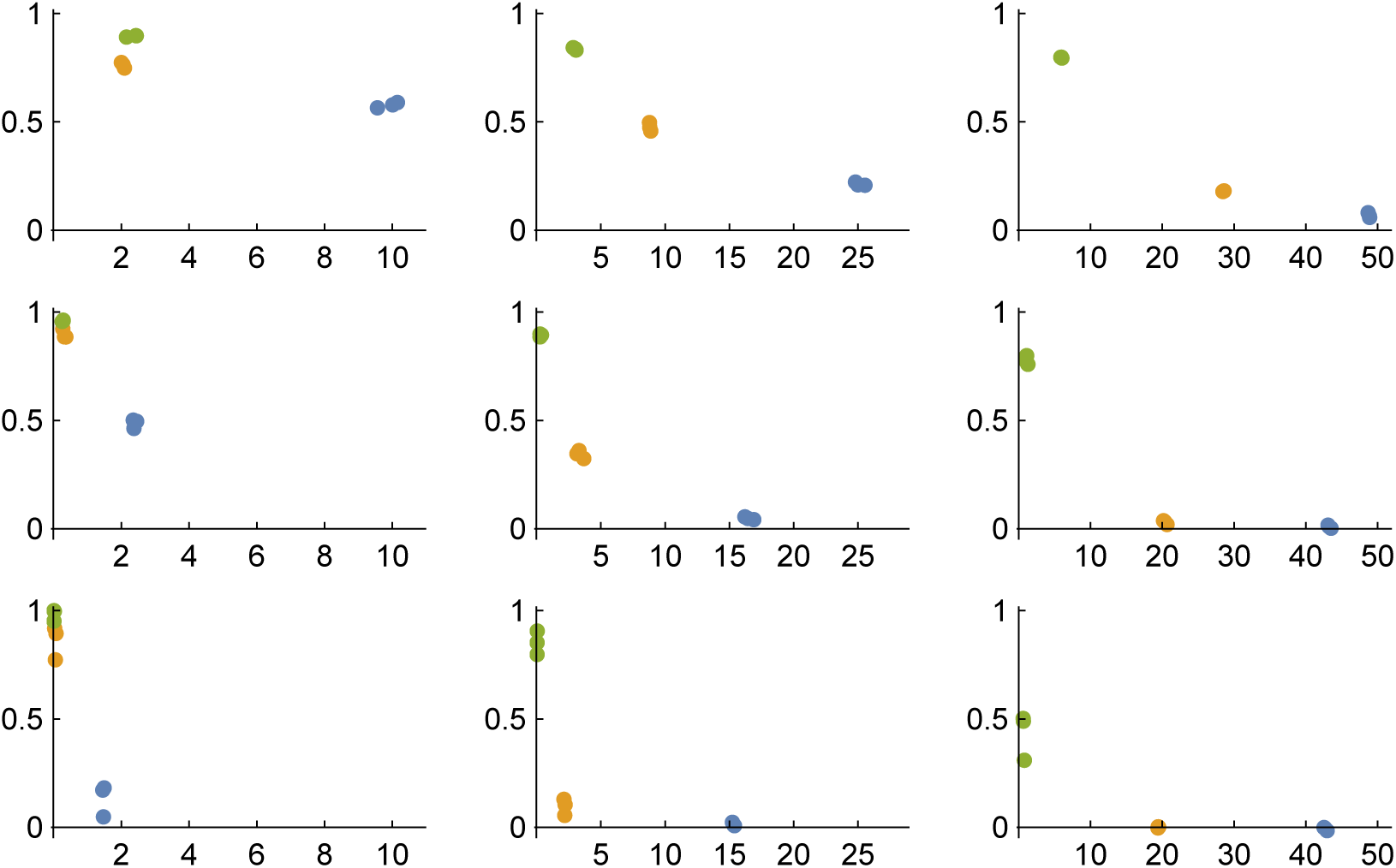
Robustness reduces disease frequency and increases the heritability of disease. The horizontal axis shows the percentage of individuals in a population with a fitness reduction of more than 10%, classified as diseased. The vertical axis shows a measure of heritability (eqn 13). The rows show rising mutation rates from bottom to top of 10^*-*4^, 10^*-*3^, 10^*-*2^. The columns show increasing intensity of selection from left to right with *σ* ^2^ *=* 10^0^, 10^*-*1^, 10^*-*2^. The colored points show increasing robustness from open loop (blue) to single feedback loop (gold) to double feedback loop (green). Each panel shows three replicate points for each colored loop type. In some panels, the three points overlap and appear as one or two. Data from Experiment D, as described in the Supplementary Information.

The vertical axes measure the tendency of disease caused by inherited genes, a measure of heritability. The figure follows the predicted pattern: increased robustness reduces the frequency of disease and raises the heritability of disease. The rows show increasing mutation rate from bottom to top. More mutation raises the frequency of diseased individuals and increases both genetic variability and the heritability of disease.

The columns show increasingly intense natural selection from left to right. Greater selection intensity means that the same phenotypic deviation in performance from the optimum yields a greater reduction in fitness. As selection intensity rises, more individuals fall into the disease category. Heritability declines because greater selection intensity reduces genetic variability and increases the relative contribution of stochastic perturbations that arise independently of genotype.

I measured heritability by recalculating fitness 100 times for each individual that fell into the disease category. Each fitness recalculation yields a different result because of stochastic perturbations. In particular, the intrinsic process parameter *α* varies randomly, and each controller parameter has a stochastic component of expression, with the amount of stochastic variability for each parameter determined by a separate genetic locus.

I scored each fitness recalculation for whether it fell within the disease category of a 10% or greater reduction in fitness from the maximum. The value of *ψ* is the frequency of disease when aggregating over all fitness recalculations for all individuals initially falling into the disease class.

I defined a heritability measure as

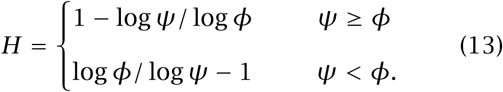

A value of *H =* 0 means that *ψ = Φ*, the frequency of recalculated disease, *ψ*, was equal to the initial frequency of disease, *Φ*. When equality occurs, recalculation of fitness for a randomly chosen individual who is initially in the disease class shows no greater tendency to fall again into the disease class than a randomly chosen individual from the entire population. Thus, *H =* 0 means that disease is, on average, not associated with inherited genetic predisposition.

A value of *H =* 1 means that *ψ =* 1, with all of the recalculated fitnesses for diseased individuals falling into the disease category. Perfect repeatability of disease upon recalculation means that inherited genetic predisposition completely determines disease.

## Conclusions

I highlight key results and promising directions for future work.

### Layering of multiple robustness mechanisms greatly improves system performance

In Fig. 5, robustness increases along the horizontal axis from an open loop (O) with no error correction to a single (S) error-correcting feedback loop to a double (D) error-correcting feedback loop.

The single loop performs better than the open loop. However, the single loop still suffers a degradation in performance when significant perturbations influence intrinsic system dynamics. By contrast, the double layering of error correction upon error correction compensates almost completely for significant perturbations to intrinsic system dynamics.

The layering of robust error-correcting mechanisms greatly boosts performance. Evolutionary dynamics may often favor such layering of error-correcting regulatory control.

### Greater robustness increases both genetic variability and stochasticity of trait expression

The top row of Fig. 7 shows the genetic variation of five different parameter values that control the system’s dynamic response to input. The center of each plot is near the optimal value. The flatter the curve, the more widely variable the genetic values for the trait. Variability rises significantly as robustness increases from an open loop (blue) to a single feedback loop (gold) to a double feedback loop (green).

The bottom row of Fig. 7 shows the inherited genetic value that determines the stochastic fluctuations in trait expression. The flatter the curve, the more high values there are in the population for stochasticity of expression. Stochasticity rises significantly as robustness increases from open loop (blue) to single feedback (gold) to double feedback (green).

### Theory makes strong comparative predictions about the relative variability of different system components

Variability in some components has relatively little effect on robust system performance, whereas variability in other components has relatively stronger effects on performance. Thus, the theory provides a fundamental basis for predicting the relative genetic variability among components and the relative stochasticity of expression.

Figure 2 shows how variation in individual components influences fitness. A broad flat curve means that large deviations in a component have small effects on fitness. For example, component *q*_2_ in the top row changes from a very narrow, sensitive curve in an open loop (blue) to a broader, less sensitive curve in a single feedback loop (gold). In the third row, the *q*_2_ panel compares the same single feedback loop (gold) with a double feedback loop (green). The double loop is significantly less sensitive than the single loop. By this theory, one expects that each additional robustness layer significantly increases the variability of *q*_2_.

By contrast, the change in sensitivity with robustness is much less for *q*_0_. Thus, the associated change in variability with robustness should be relatively lower for *q*_0_ than for *q*_2_. Those theory curves were obtained by analyzing the sensitivity of system fitness to variations in each component, holding all other components at their optimum value. In real systems, all components will vary simultaneously.

The computer simulation output in Fig. 7 supports the prediction that the variability of *q*_2_ (fifth column) changes more with increasing robustness than the variability of *q*_0_ (third column).

### Robustness reduces the frequency of disease

I measured disease intensity by the fraction of the population with fitness reduced by at least *X*%. In the simulation analysis, I used 10%. Robustness reduced disease frequency, as shown in Fig. 8. The horizontal axis of each panel is the percentage of individuals classified as diseased. As robustness increased from open loop (blue) to single feedback loop (gold) to double feedback loop (green), disease frequency declined.

It may seem obvious that robustness should decrease disease frequency. However, the evolutionary dynamics of robustness in relation to disease can be complex. As system robustness increases, fitness improves. But simultaneously, as system robustness increases, components tend to decay in performance, which reduces fitness (Frank, 2004, 2007b; Lynch, 2012).

The net effect of robustness on system performance depends on the balance between the initial gain in system performance and the decay in system performance that follows as components decay. In this case, the net effect was a significant decline in disease as robustness increased. At present, no general theory clarifies how the opposing forces balance at equilibrium and the consequences for disease frequency. I suspect there is some economic marginal law of gains and losses in performance that explains the balance at equilibrium.

### Robustness increases the heritability of disease and the heritability of fitness

A feedback system corrects error. Good error correction requires a reasonably accurate estimate of errors and a mechanism to respond appropriately to error estimates. Thus, robust systems may be more sensitive to inherited changes that alter intrinsic error correction than to stochastic perturbations that can be compensated by error-correcting feedback.

In other words, greater robustness may lead to a higher heritability of performance, because the genetically determined error-correction system dominates the performance of highly robust systems. To analyze that prediction in the simulations, I measured the heritability of disease. For each individual that fell below the cutoff of a 10% reduction in fitness from the optimum, I recalculated fitness 100 times. Each recalculation included random perturbations to the system dynamics.

I measured heritability by the tendency for an individual initially classified as diseased to repeatedly fall into the disease class. The higher the repeatability, the more strongly disease is determined by inherited genotype rather than stochastic perturbations. Figure 8 shows that increasing robustness strongly enhances the heritability of disease.

Perhaps intrinsic robustness partially explains the puzzlingly high heritability of many diseases (Frank, 2007a; Manolio et al., 2009; Eichler et al., 2010). Also, fitness itself is often highly heritable. That high heritability for fitness has been considered a puzzle because natural selection should, in principle, rapidly remove genetic variability for fitness (Mousseau & Roff, 1987). Robustness may explain the high heritability of fitness.

### Promising directions for future work

Each additional layer of robustness causes the lower-level components to decay. How are system design dynamics influenced by this inevitable system layering and component decay? How does layering and system design by evolutionary processes differ from design by human designers?

It would seem that design dynamics would favor substituting a cheaper, lower-performing component when a system becomes protected by an additional robustness mechanism (Frank, 2007b). How important are such economic considerations in understanding design dynamics?

I focused on the simplest type of error correction. How do the various advanced types of robustness mechanisms alter design dynamics? Control theory provides tools to analyze adaptive (learning) robustness mechanisms and other mechanisms such as model predictive control (Frank, 2018a). Can we develop general understanding of the various types of robustness and their consequences for evolutionary dynamics?

The theory predicts that different system components will accumulate different amounts of variability in response to robustness. Such predictions provide a theory for the relative levels of genetic variability and stochasticity of trait expression. Modern technology provides much data about genetic variability and single-cell stochasticity of gene expression. In principle, it should be possible to test the theory against those data.

However, an open challenge remains to connect abstract notions of error-correcting dynamics to individual genes and to particular molecules within complex biochemical networks of reactions (Frank, 2018b). Making that connection between function and mechanism remains an essential challenge for understanding the design of biological systems.

## Acknowledgments

National Science Foundation grant DEB–1251035 and the Donald Bren Foundation support my research. I completed this work while on sabbatical in the Theoretical Biology group of the Institute for Integrative Biology at ETH Zürich.

## Supplemental files

The Supplementary Information, the Mathematica code used to analyze the simulation data and produce the figures, and the raw simulation data are available at https://doi.org/10.5281/zenodo. 1405960. The C++ source code is available at https://doi.org/10.5281/zenodo.1405968.

## References

Åström, K. J. & Murray, R. M. (2008). Feedback Systems: An Introduction for Scientists and Engineers (Version v2.11a ed.). Princeton, NJ: Princeton University Press.

de Visser, J., Hermisson, J., Wagner, G. P., Meyers, L. A., Bagheri-Chaichian, H., Blanchard, J. L., Chao, L., Cheverud, J. M., Elena, S. F., Fontana, W., Gibson, G., Hansen, T. F., Krakauer, D., Lewontin, R. C., Ofria, C., Rice, S. H., von Dassow, G., Wagner, A., & Whitlock, M. C. (2003). Perspective: evolution and detection of genetic robustness. Evolution, 57, 1959–1972.

Dorf, R. C. & Bishop, R. H. (2016). Modern Control Systems (13th ed.). Santa Monica, California: Pearson.

Eichler, E. E., Flint, J., Gibson, G., Kong, A., Leal, S. M., Moore, J. H., & Nadeau, J. H. (2010). Missing heritability and strategies for finding the underlying causes of complex disease. Nature Reviews Genetics, 11(6), 446.

Frank, S. A. (2004). Genetic variation in cancer predisposition: mutational decay of a robust genetic control network. Proceedings of the National Academy of Sciences USA, 101, 8061–8065.

Frank, S. A. (2007a). Dynamics of Cancer: Incidence, Inheritance, and Evolution. Princeton, NJ: Princeton University Press.

Frank, S. A. (2007b). Maladaptation and the paradox of robustness in evolution. PLoS ONE, 2, e1021.

Frank, S. A. (2013). Evolution of robustness and cellular stochasticity of gene expression. PLoS Biology, 11, e1001578.

Frank, S. A. (2018a). Control Theory Tutorial: Basic Concepts Illustrated by Software Examples. Cham, Switzerland: Springer.

Frank, S. A. (2018b). Evolutionary design of regulatory control. I. A robust control theory analysis of tradeoffs. bioRxiv https://doi.org/10.1101/332999.

Lynch, M. (2012). Evolutionary layering and the limits to cellular perfection. Proceedings of National Academy Sciences USA, 109, 18851–18856.

Manolio, T. A., Collins, F. S., Cox, N. J., Goldstein, D. B., Hindorff, L. A., Hunter, D. J., McCarthy, M. I., Ramos, E. M., Cardon, L. R., Chakravarti, A., et al. (2009). Finding the missing heritability of complex diseases. Nature, 461(7265), 747.

Mousseau, T. A. & Roff, D. A. (1987). Natural selection and the heritability of fitness components. Heredity, 59(2), 181.

Ogata, K. (2009). Modern Control Engineering (5th ed.). New York: Prentice Hall.

Rutherford, S. L. & Lindquist, S. (1998). Hsp90 as a capacitor for morphological evolution. Nature, 396(6709), 336–342.

Wagner, A. (2013). Robustness and Evolvability in Living Systems. Princeton University Press.

